# Integrating multimodal connectivity improves prediction of individual cognitive abilities

**DOI:** 10.1101/2020.06.25.172387

**Authors:** Elvisha Dhamala, Keith W. Jamison, Abhishek Jaywant, Sarah Dennis, Amy Kuceyeski

## Abstract

How white matter pathway integrity and neural co-activation patterns in the brain relate to complex cognitive functions remains a mystery in neuroscience. Here, we integrate neuroimaging, connectomics, and machine learning approaches to explore how multimodal brain connectivity relates to cognition. Specifically, we evaluate whether integrating functional and structural connectivity improves prediction of individual crystallised and fluid abilities in 415 unrelated healthy young adults from the Human Connectome Project. Our primary results are two-fold. First, we demonstrate that integrating functional and structural information – at both a model input or output level – significantly outperforms functional or structural connectivity alone to predict individual verbal/language skills and fluid reasoning/executive function. Second, we show that distinct pairwise functional and structural connections are important for these predictions. In a secondary analysis, we find that structural connectivity derived from deterministic tractography is significantly better than structural connectivity derived from probabilistic tractography to predict individual cognitive abilities.

## Introduction

Tens of billions of neurons interconnect in the human brain. Direct and indirect structural white matter connections between these neurons facilitate the flow of functional activation between distinct brain regions. Together, these functional and structural connections give rise to human behaviour and cognition. Insight into multimodal neural correlates of cognitive abilities in the healthy brain provides an important foundation with which to delineate age-, injury-, and disease- related changes in cognitive functioning. Furthermore, a thorough understanding of specific functional and structural connections that are associated with cognition can guide the investigation of causality, and possible, the development of targeted neuromodulatory treatments for cognitive dysfunction.

Functional connectivity (FC) represents temporal dependency patterns between regional blood-oxygenation-level dependent (BOLD) activity in functional magnetic resonance imaging (fMRI) time series, and structural connectivity (SC) represents the integrity of inter-regional white matter pathways estimated from diffusion MRI (dMRI). FC and SC have individually been linked to cognitive functioning and used to predict cognitive measures (Song et al., 2009, Song et al., 2008, Pamplona et al., 2015, Klein et al., 2016, Moeller et al., 2015, Willmes et al., 2014, Matejko et al., 2013, Seeley et al., 2007, van den Heuvel et al., 2009, Zimmermann et al., 2018, He et al., 2020, Kong et al., 2018, Li et al., 2019, Liégeois et al., 2019, Bassett et al., 2011, Menon and Uddin, 2010, Uddin et al., 2011, Kelly et al., 2008). Although neural function and structure are inexorably linked, most studies analyse their contribution to behaviour independently. FC is associated with performance variability in executive control (Seeley et al., 2007) and intellectual performance (van den Heuvel et al., 2009), and can successfully predict a range of cognitive measures (He et al., 2020, Kong et al., 2018, Li et al., 2019). Moreover, morphometric similarity networks capturing neuroanatomical properties from structural and diffusion images (fractional anisotropy, mean diffusivity, magnetisation transfer, grey matter volume, surface area, cortical thickness, intrinsic curvature, mean curvature, curved index, and folding index) can explain variability in general intellectual functioning (Seidlitz et al., 2018), and structural connectivity can accurately predict fluid abilities (Zimmermann et al., 2018). FC and SC seem to independently contribute to cognition (Zimmermann et al., 2018), but no work has yet investigated whether integrating FC and SC can increase cognitive prediction accuracy above and beyond what is obtained by either modality alone. Here, we sought to integrate FC and SC (Amico and Goñi, 2018) to reveal how both modalities contribute to predicting cognition.

Most studies to date have focused on using resting-state FC to predict behavioural and cognitive measures (He et al., 2020, Kong et al., 2018, Li et al., 2019). Kong et al. (Kong et al., 2018) used spatial topography of cortical functional networks to predict behaviour. Li et al. (Li et al., 2019) found global signal regressed resting-state FC improves behavioural prediction. He at al. (He et al., 2020) showed that machine learning and deep learning methods are equally effective in predicting behavioural, cognitive, and demographic measures from resting-state FC. However, these studies have not addressed whether SC can provide additional explained variance in cognition.

In the most similar study to date, Zimmerman et al. (Zimmermann et al., 2018) studied multi-dimensional connectome-cognition relationships in 609 genetically unrelated subjects from the Human Connectome Project (Van Essen et al., 2013). They generated three main components from eleven cognitive measures and used partial least squares analyses to identify four latent variables that describe the connectome-cognition relationships: two captured FC-cognition associations and two captured SC-cognition associations (Zimmermann et al., 2018). They found functional and structural connections uniquely map onto cognitive functions, including working memory, executive function, cognitive flexibility, processing speed, fluid intelligence, episodic memory, and attention/inhibitory control (Zimmermann et al., 2018). They identified a large set of distributed interhemispheric functional connections spanning bilateral frontal, parietal, temporal, and subcortical regions, and a limited set of short-range intrahemispheric structural connections within bilateral parietal, temporal and subcortical regions that distinctively map onto cognitive function (Zimmermann et al., 2018). While this study addressed the relationship between SC and cognition, it analysed the functional and structural relationships to cognition independently.

Amico and Goñi (Amico and Goñi, 2018) have simultaneously studied changes in functional and structural connectivity patterns across tasks and resting-state. Their results showed that combining functional and structural connectivity into a “hybrid” connectivity, they were able to extract meaningful information and capture individual differences (Amico and Goñi, 2018). However, the relationship between this “hybrid” connectivity and cognitive functioning has not yet been explored.

White matter pathways comprise neural circuits involving cortical and subcortical brain regions. Tractography can be used to estimate these pathways from dMRI. Two main classes of tractography algorithms, deterministic and probabilistic, differ in how they sample fibre directions for streamline propagation. Whether structural connectivity derived from deterministic or probabilistic tractography is better at predicting individual cognitive measures has not yet been established.

Here, we study whether integrating SC and FC improves cognitive predictions in a subset of 415 healthy young adults from the Human Connectome Project (Van Essen et al., 2013) dataset. The primary goals of this study are twofold. First, we evaluate whether integrating functional and structural connectivity, at either a model input or output level, improves the prediction of individual crystallised and fluid cognitive abilities when compared to functional and structural connectivity alone. Second, we quantify the pairwise functional and structural connections that contribute the most to the prediction of crystallised and fluid abilities. As a secondary analysis, we compare whether structural connectivity derived from deterministic or probabilistic tractography is superior at predicting individual cognitive abilities.

## Methods

Our experimental workflow is shown in Figure 1. The data that support the findings of this study are openly available as part of the Human Connectome Project at https://www.humanconnectome.org/study/hcp-young-adult/document/1200-subjects-data-release (Van Essen et al., 2013). The codes generated during this study are available on GitHub (https://github.com/elvisha/CognitivePredictions).

**Figure 1:**
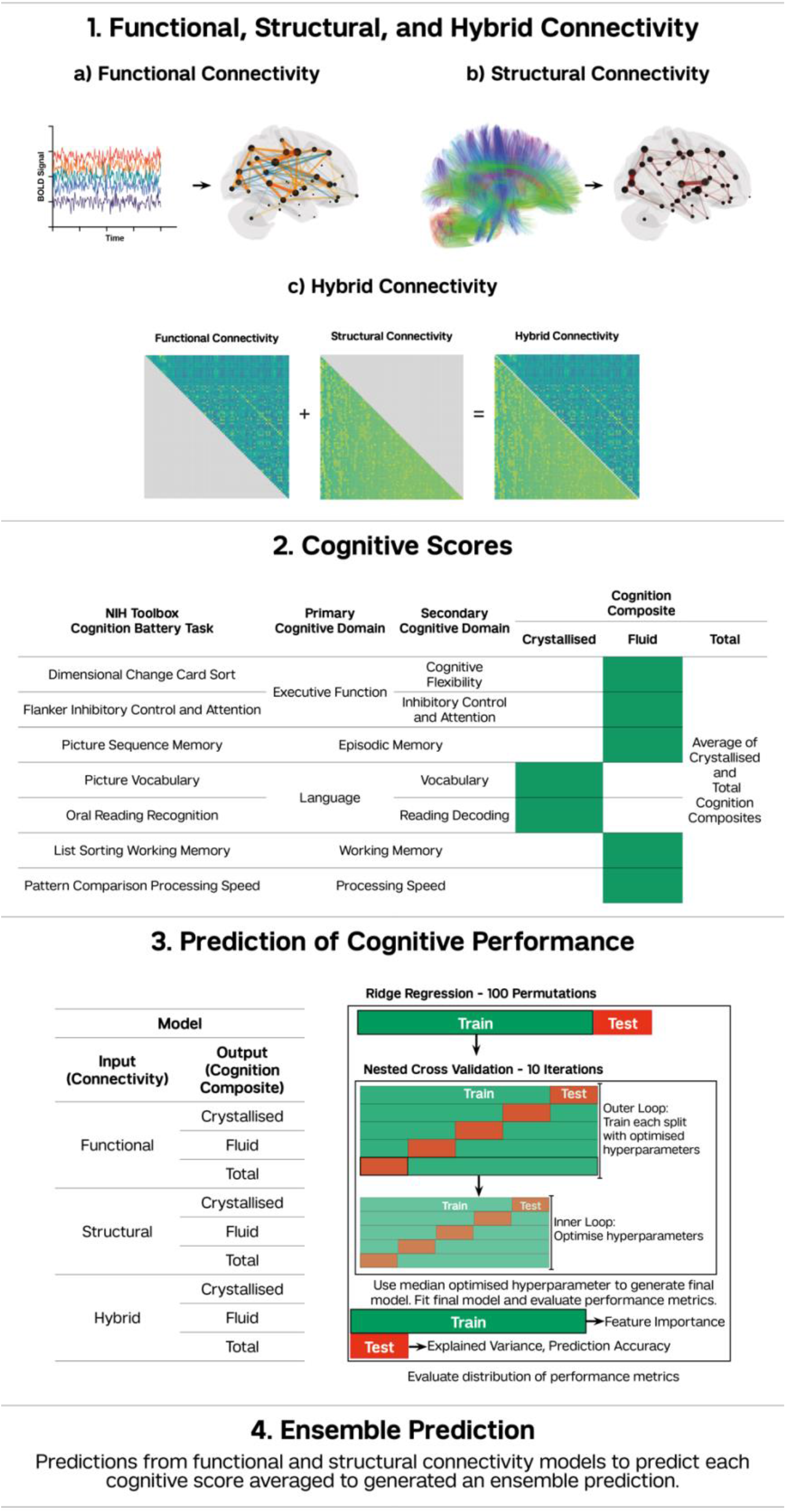
Workflow of Experiment. First, we generated the functional, structural and hybrid connectivity matrices (1). We derived functional connectivity using zero-lag Pearson correlation of global signal regressed blood-oxygen-level-dependent (BOLD) functional MRI time series (1a) and Fisher’s z-transformed the upper triangular matrix. We derived structural connectivity using deterministic and probabilistic tractography from diffusion weighted MRI (1b) and Gaussian resampled the lower triangular matrix. We concatenated the upper triangular Fisher’s z-transformed functional connectivity and lower triangular Gaussian resampled structural connectivity matrices to generate hybrid connectivity (1c). Second, we obtained cognitive scores for all subjects (2). The NIH Toolbox Cognition Battery assesses five cognitive domains using seven tests. The Crystallised Cognition Composite reflects language (vocabulary, reading decoding). The Fluid Cognition Composite reflects executive function (cognitive flexibility, inhibitory control and attention), episodic memory, working memory, and processing speed. The Total Cognition Composite is the average of Crystallised and Fluid Cognition Composite scores. Third, we predicted each of the cognitive scores using each of the connectivity matrices (3) via linear ridge regression models. We first split the data into train (80%) and test (20%) subsets. We then performed ten iterations of nested cross validation with five-fold inner and five-fold outer loops on the train subset to optimise hyperparameters; we fit the final model using the median optimised hyperparameter. We performed 100 permutations of each model to generate a distribution of performance metrics and feature importances using unique train/test splits. Fourth, we averaged predictions from the functional and structural connectivity models to generate an ensemble prediction for each test subject’s Crystallised, Fluid, and Total Cognition Composites.

### Dataset

We used publicly-available high resolution, preprocessed magnetic resonance imaging (MRI) data from the Human Connectome Project – Young Adult S1200 (Van Essen et al., 2013) in this study. HCP MRI data were acquired on a Siemens Skyra 3T scanner at Washington University in St. Louis. HCP scanning included T1-weighted and T2-weighted anatomical images (0.7mm isotropic), functional MRI (2.0mm isotropic, TR/TE = 720/33.1ms, 8x multiband acceleration), and diffusion MRI (1.25mm isotropic, TR/TE = 5520/89.5ms, 3x multiband acceleration, b=1000,2000,3000, 90 directions/shell). Functional and diffusion MRI were collected with both left-right and right-left phase encoding. We examined resting-state functional MRI (rfMRI) time series and diffusion MRI (dMRI) from 415 unrelated healthy adults (213 males; ages 22-37). This subset of the HCP dataset were those subjects that had four complete rfMRI runs, a dMRI scan and crystallised and fluid cognitive scores.

### Parcellation

As part of the HCP preprocessing workflow (Glasser et al., 2013), FreeSurfer’s recon-all pipeline (Dale et al., 1999, Fischl et al., 2001, Fischl et al., 2008, Fischl et al., 2002, Fischl et al., 1999a, Fischl et al., 1999b, Segonne et al., 2005) was optimised for the high-resolution HCP anatomical data. The 68 region Desikan-Killiany gyral atlas (aparc.annot, 34 cortical regions per hemisphere) was combined with the 18 bilateral subcortical structures (aseg.mgz, excluding brainstem) to produce an 86 region whole brain anatomical parcellation for each subject (Desikan et al., 2006, Fischl et al., 2002). We parcellated the brain into these 86 regions to generate both the functional and structural connectomes as described below to maintain consistency and enable comparisons between the two.

### Functional Connectome (FC) Extraction

Each subject underwent four gradient-echo EPI rfMRI runs of ~15 min each over two sessions. The data consisted of 1200 volumes per rfMRI for a total of 4800 volumes for each subject over the four runs. The minimal preprocessing pipeline performed by the HCP consortium included motion and distortion correction, registration to subject anatomy and standard MNI space, and automated removal of noise artefacts by independent components analysis(Glasser et al., 2013, Griffanti et al., 2014, Salimi-Khorshidi et al., 2014). We regressed the global signal and its temporal derivative from each rfMRI time series and computed the zero lag Pearson correlation to derive the FC for each scan. For each subject, we averaged FC across the four scans to get a single mean FC, which we then Fisher’s z-transformed. We used the vectorised upper triangular of this FC to predict cognition.

### Structural Connectome (SC) Extraction

The HCP minimally preprocessed diffusion data have been processed to correct for motion, EPI and eddy-current distortion, and registered to subject T1 anatomy (Glasser et al., 2013). We then used MRtrix3 to estimate a voxel-wise multi-shell, multi-tissue constrained spherical deconvolution (CSD) model and then compute whole brain tractography for each HCP subject (Jeurissen et al., 2014). We computed separate whole-brain tractograms using both probabilistic (iFOD2 (Tournier et al., 2010) with anatomically constrained tractography – ACT (Smith et al., 2012)) and deterministic (SD_STREAM (Tournier et al., 2012)) tractography algorithms. Each method produced 5 million streamlines per subject, using dynamic seeding, and computed streamline weights to reduce known biases in tractography algorithms and better match the whole brain weighted tractogram to diffusion properties of the observed data (SIFT2, (Smith et al., 2015)). We parcellated the tractograms to produce ROI-volume normalized pairwise SC matrices, where each pairwise connection is the sum of the SIFT2 weights of streamlines connecting those regions, divided by the sum of the grey matter volume of those regions. We generated two SC matrices for each subject: one using deterministic tractography and another using probabilistic tractography. We resampled SC matrices independently to a Gaussian distribution (Honey et al., 2009): given *N* raw data values x_1_,…,x_N_, *N* random samples r_1_,…,r_N_, from a Gaussian distribution with a mean of 0.5 and a standard deviation of 0.1 were generated. We replaced the smallest raw data value with the smallest randomly sampled value and repeated until all raw data values were replaced. This produced a set of *N* resampled data values with a Gaussian distribution. We used the vectorised lower triangular of this Gaussian resampled SC to predict cognition.

### Hybrid Connectome (HC)

We concatenated the upper triangular of the functional and the lower triangular of the structural connectivity matrices to generate HC (Amico and Goñi, 2018). We used the vectorised HC to predict cognition.

### Cognition

The NIH Toolbox Cognition Battery is an extensively validated battery of neuropsychological tasks (Carlozzi et al., 2017, Gershon et al., 2013, Heaton et al., 2014, Mungas et al., 2014, Tulsky et al., 2017, Weintraub et al., 2013, Weintraub et al., 2014, Zelazo et al., 2014) that assesses five cognitive domains: language, executive function, episodic memory, processing speed, and working memory through seven individual test instruments (Heaton et al., 2014). The specific tasks include Dimensional Change Card Sort Test (executive function – cognitive flexibility), Flanker Inhibitory Control and Attention Test (executive function – inhibitory control and attention), Picture Sequence Memory Test (episodic memory), Picture Vocabulary Test (language – vocabulary), Oral Reading Recognition Test (language – reading decoding), List Sorting Working Memory Test (working memory), and Pattern Comparison Processing Speed Test (processing speed) (Heaton et al., 2014). Three composite scores are derived from participants’ scores on the NIH Toolbox Cognitive Battery tasks: Crystallised Cognition Composite, Fluid Cognition Composite, and Total Cognition Composite (Heaton et al., 2014). The Crystallised Cognition Composite comprises the Picture Vocabulary and Oral Reading Recognition tests and assesses language and verbal skills. The Fluid Cognition Composite comprises scores on the Dimensional Change Card Sort, Flanker Inhibitory Control and Attention, Picture Sequence Memory, List Sorting Working Memory, and Pattern Comparison Processing Speed tests. It is a composite that broadly assesses processing speed, memory, and executive functioning. The Total Cognition Composite is the average of the Crystallised and Fluid Cognition Composites. We used the Crystallised, Fluid, and Total Cognition Composites in this study, rather than the individual scores from the tasks, because they are likely to have a higher signal-to-noise ratio. Composite scores also tend to be more reliable/stable and are less susceptible to variability in individual tasks (Heaton et al., 2014). Lastly, by using the composite scores, we greatly reduce the number of models that need to be trained, thus reducing the number of multiple comparison.

### Prediction of Cognitive Performance

We used three distinct inputs (FC, SC, and HC) to predict three distinct outputs (crystallised, fluid, and total cognition): a separate machine learning model was trained for each input/output combination. To evaluate whether SC derived from deterministic or probabilistic tractography is superior at predicting cognition, we trained two separate models for SC-based predictions. For each model, we split the data into training (80%) and testing (20%) splits. We fit a linear ridge regression model on Scikit-learn (Pedregosa et al., 2011) using the training subset and tuned the regularisation parameter with ten iterations of nested cross validation with five-fold inner and outer loops. We optimised the regularisation parameter in the inner loop and then used it to train each split in the outer loop. We took the median over the optimised hyperparameters from the ten iterations to generate a single final model. We trained this model and extracted feature weights from the training set and evaluated the model’s explained variance and prediction accuracy from the test set. We quantify prediction accuracy as the Pearson correlation between the true and predicted values (Li et al., 2019). We performed one hundred permutations of each model to generate a distribution of performance metrics while maintaining the distinct train/test splits consistent for all prediction models. We also trained Elastic Net, LASSO, and kernel ridge regression models to predict cognition, but results were inferior to those from linear ridge regression models. Hence, we specifically report results from linear ridge regression. Finally, we generated an Ensemble Prediction (EP) for each test subject’s Crystallised, Fluid, and Total Cognition Composite by averaging the predictions from models trained independently on FC and SC (Khosla et al., 2019). For each cognitive score, we evaluated hold-out performance differences between the FC, SC, HC and EP models using a Kruskal-Wallis test. We corrected p-values for multiple comparisons using the Benjamini-Hochberg procedure (Benjamini and Hochberg, 1995) to decrease the false discovery rate. To quantify the effect of tractography algorithm choice on the accuracy of cognitive prediction, we evaluated the difference in model performance metrics for all three cognitive scores using SC derived from deterministic versus probabilistic tractography using a Mann-Whitney *U* test.

### Feature Importance

We averaged feature weights obtained over the 100 permutations of the linear ridge regression models to get a mean feature weight. Feature weights from HC models were separated into their functional and structural components. We rescaled the feature weights while maintaining their signs (positive and negative) to generate pairwise feature importance. We computed regional positive and negative feature importance by taking the sum over that region’s pairwise positive and negative feature importances. We evaluated the correlations between pairwise feature weights obtained from FC, SC, and HC models to predict crystallised, fluid, and total cognitive scores. We corrected p-values corresponding to the correlations for multiple comparisons using the Benjamini-Hochberg procedure (Benjamini and Hochberg, 1995) to decrease the false discovery rate.

Further information and requests for resources should be directed to and will be fulfilled by the Lead Contact, Elvisha Dhamala.

## Results

### Prediction of Cognitive Performance

FC, SC, HC, and EP explain 11.3% (prediction accuracy, r=0.34), 8.6% (r=0.31), 14.6% (r=0.39), and 12.5% (r=0.40) of the variance in crystallised cognition, respectively. FC, SC, HC, and EP explain 6.2% (r=0.25), 5.5% (r=0.24), 8.3% (r=0.29), and 7.9% (r=0.32) of the variance in fluid cognition, respectively. FC, SC, HC, and EP explain 11.3% (r=0.34), 11.5% (r=0.34), 14.7% (r=0.39), and 13.9% (r=0.41) of the variance in total cognition, respectively. We evaluated differences in explained variance and prediction accuracy between the models using FC, SC, HC, and EP to predict each of the cognitive scores. In terms of prediction accuracy, HC and EP significantly (p<0.05) outperform FC and SC alone to predict crystallised, fluid, and total cognitive abilities. In terms of explained variance: FC and EP significantly (p<0.05) outperform SC, and HC significantly (p<0.05) outperforms FC and SC alone to predict crystallised abilities; HC significantly (p<0.05) outperforms FC and SC alone, and EP significantly (p<0.05) outperforms SC alone to predict fluid abilities; HC and EP significantly (p<0.05) outperform FC and SC alone to predict total cognitive abilities. HC and EP models perform comparably (p>0.05) to predict crystallised, fluid, and total cognition. Figure 2 shows explained variance and prediction accuracy violin plots for models using FC, SC, HC, and EP to predict crystallised, fluid, and total cognition.

**Figure 2:**
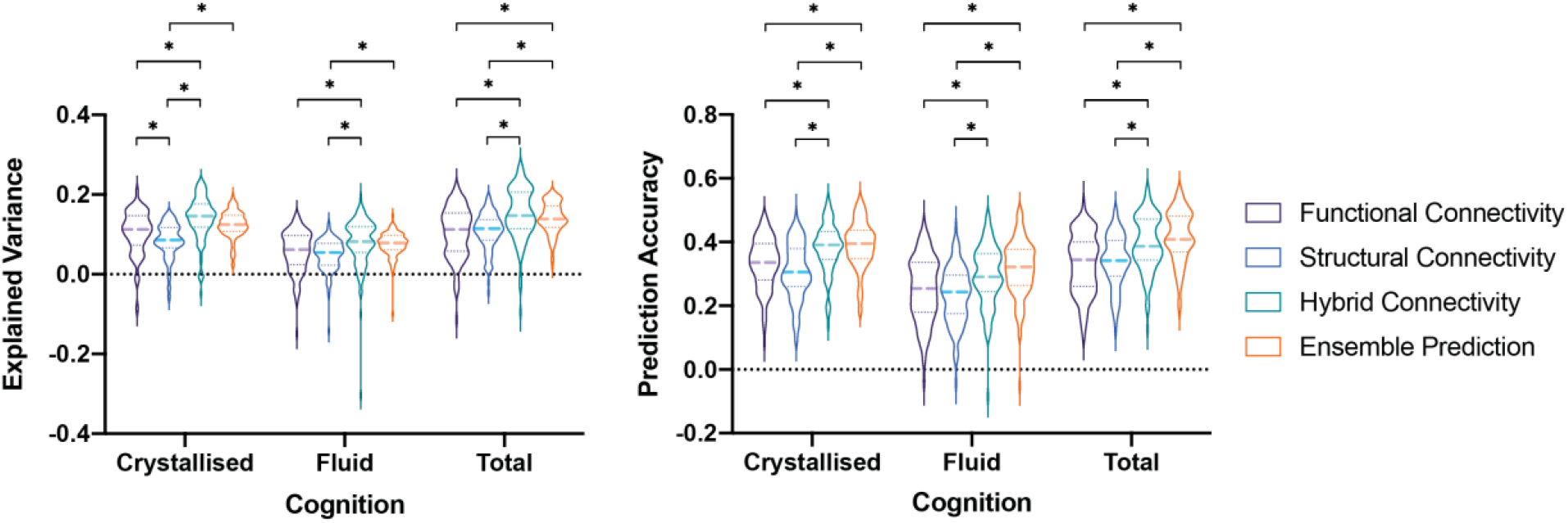
Model Performance Metrics from Prediction of Cognitive Performance. Explained variance (left) and prediction accuracy (right) violin plots for models using functional connectivity, structural connectivity, hybrid connectivity, and ensemble predictions to predict crystallised, fluid, and total cognition. Structural connectivity, hybrid connectivity, and ensemble prediction models here used structural connectivity derived from deterministic tractography. Solid lines indicate the distribution of values, dashed lines indicate the median, and dotted lines indicate the interquartile range. Significant differences in model performance between the four connectivity inputs were evaluated for each of the cognitive scores using a Kruskal-Wallis test. * indicates significant differences (p < 0.05) after Benjamini-Hochberg correction for multiple comparisons.

### Feature Importance and Regional Importance

Distributed pairwise functional and structural connections are important to predict individual cognitive scores. Fronto-temporal and cortico-subcortical functional connections and structural connections between cortical regions and the left caudate are important in predicting crystallised scores. In contrast, the feature importance scores for fluid cognition show an even more distributed set of cortical-subcortical functional connections and an even larger emphasis on structural connections out of the left caudate. To predict individual total cognition, a combination of the connections important to predict crystallised and fluid cognition are important. Pairwise functional and structural connections important to predict individual crystallised, fluid, and total cognition abilities are shown in Figure 3.

**Figure 3:**
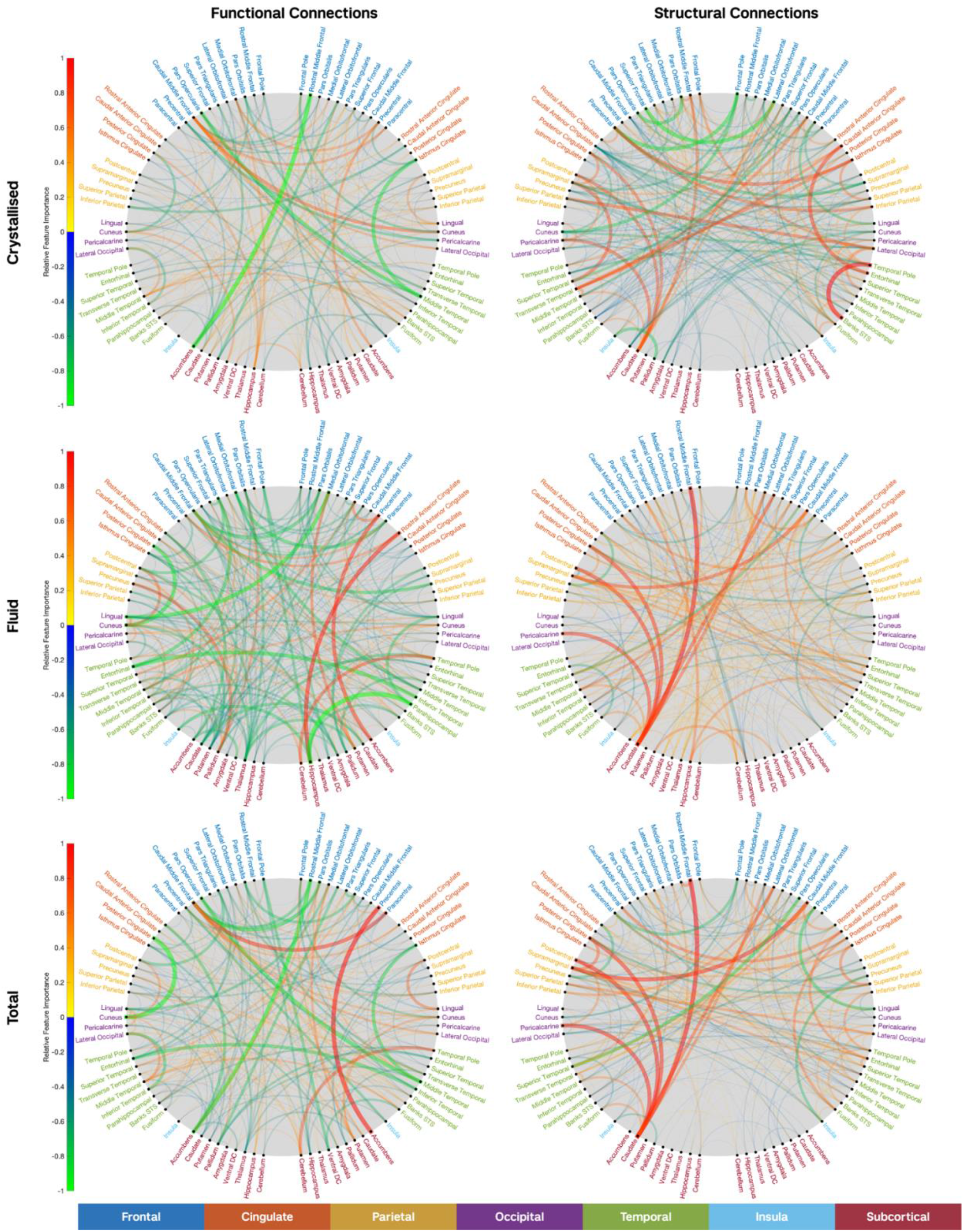
Pairwise Feature Importance. Pairwise feature importance from models using functional connectivity (left) and structural connectivity (right) to predict Crystallised (top), Fluid (middle), and Total (bottom) Cognition Composites. The top 20% most important pairwise connections are shown. Warm colours denote positive feature weights and cool colours denote negative feature weights. Nodes along the graph are organised based on lobes as per the colour map, and the right side of the graph corresponds to the right side of the brain.

Distinct functional and structural region-pair connections are important for predicting each of the cognitive scores. That is, for each cognitive score, there exists no correlation (p>0.05) between pairwise feature importances from models based on functional or structural connectivity. However, there exist significant correlations (p<0.05) between pairwise feature importances from models using functional connectivity to predict each of the three cognitive scores and structural connectivity to predict each of the three cognitive scores. Pairwise feature importances of functional connections from the functional and hybrid connectivity models and structural connections from the structural and hybrid connectivity models are significantly correlated (p<0.05) for each cognitive score. Correlations between pairwise feature importances are shown in Figure 4.

**Figure 4:**
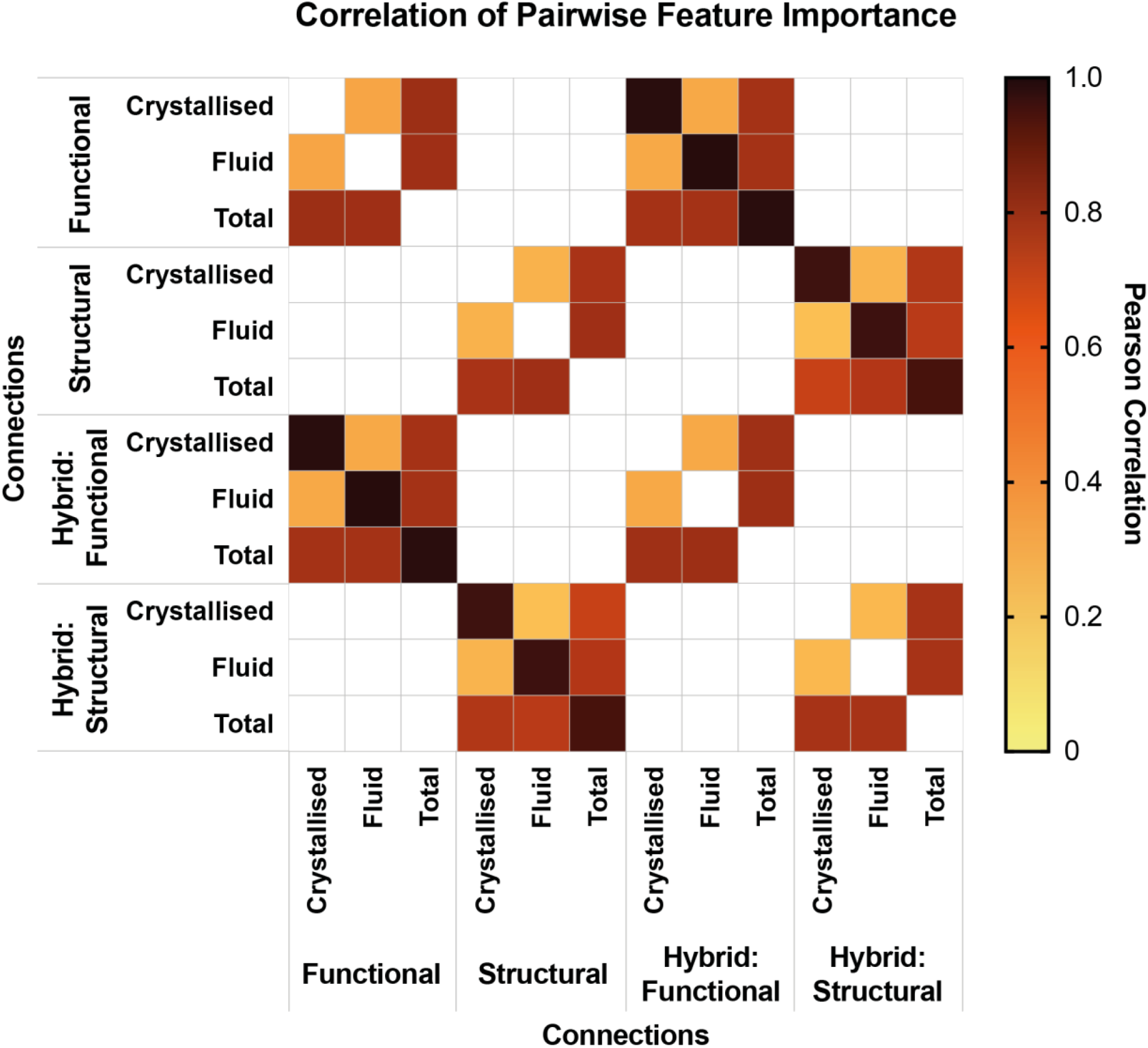
Correlation of Pairwise Feature Importance. Correlation of pairwise functional and structural feature weights extracted from models using functional, structural, and hybrid connectivity to predict Crystallised, Fluid, and Total Cognition Composites. Pearson correlation for all significant correlations (p<0.05) after Benjamini-Hochberg correction for multiple comparisons are shown. Hotter colours indicate stronger correlation. Non-significant correlations are shown in white.

Similar to the feature importance of the region-pair connections, different regions have overall importance in predicting individual cognitive scores. For each cognitive score, there exists no correlation (p>0.05) between regional feature importances (calculated as sum of absolute pairwise connection feature importance for each region) from models using FC or SC. The SC of a large, distributed set of cortical regions in the bilateral frontal, parietal, temporal, and occipital cortices are important in predicting individual crystallised and fluid abilities; the FC of a smaller set of regions in the bilateral frontal, left temporal, and left subcortical cortices are important, see Figure 5. Regional positive and negative feature importance is shown in Supplementary Figure 1.

**Figure 5:**
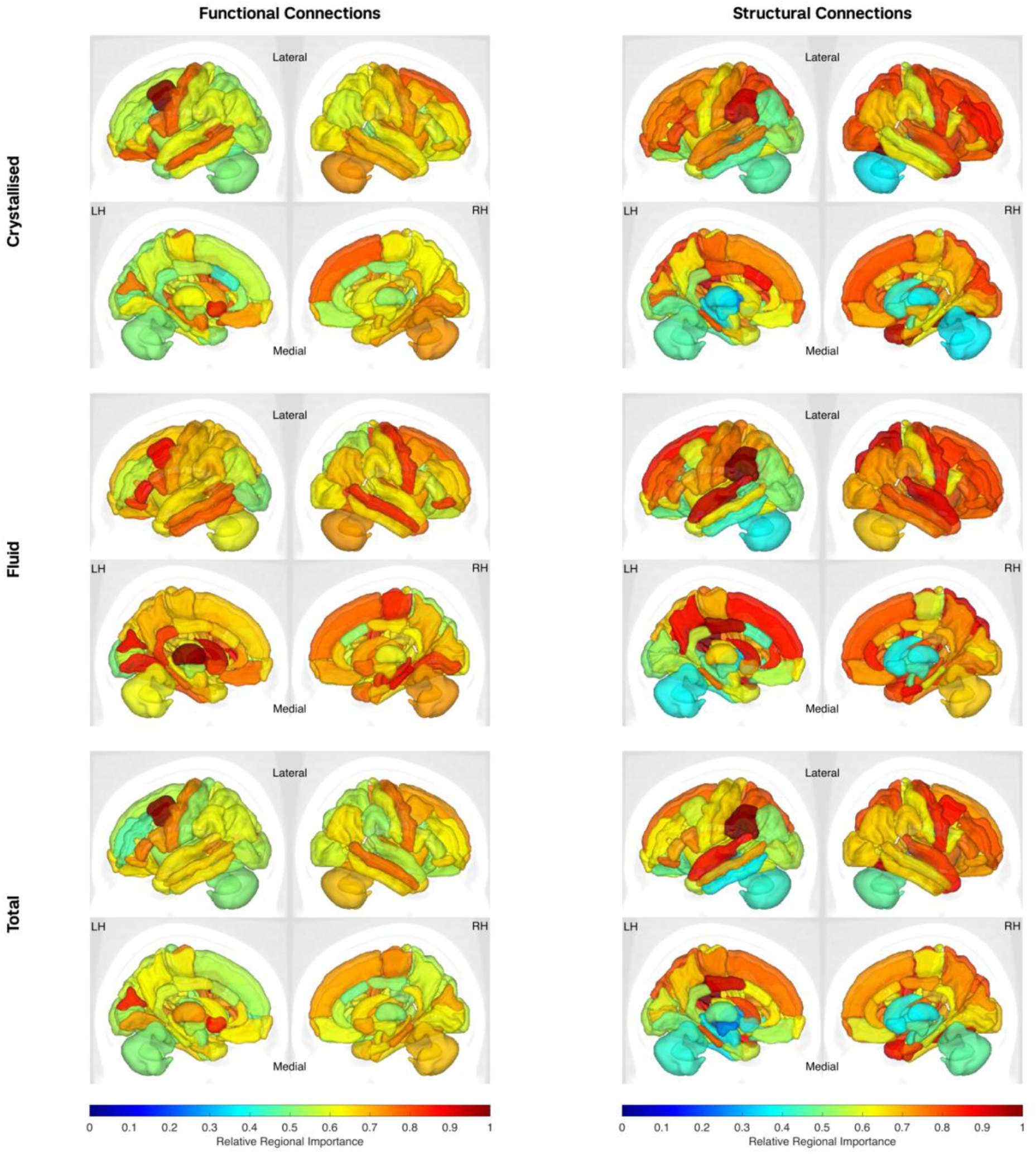
Regional Feature Importance. Regional feature importance from models using functional connectivity (left) and structural connectivity (right) to predict Crystallised (top), Fluid (middle), and Total (bottom) Cognition Composites. Relative regional importance is shown as per the colourmap. Lateral and medial views of the right (RH) and left (LH) hemispheres are shown.

### Deterministic versus Probabilistic Tractography

SC derived from deterministic tractography explains 8.6% (prediction accuracy, r=0.31), 5.5% (r=0.25), and 11.5% (r=0.34) of the variance in crystallised, fluid, and total cognition, respectively, while SC derived from probabilistic tractography explains 5.0% (r=0.23), 3.2% (r=0.18), and 6.2% (r=0.26), respectively. SC derived from deterministic tractography significantly outperforms SC derived from probabilistic tractography when predicting crystallised, fluid, and total cognition in terms of both explained variance and prediction accuracy. Explained variance and prediction accuracy violin plots to predict crystallised, fluid, and total cognition using SC derived from deterministic and probabilistic tractography are shown in Figure 6.

**Figure 6:**
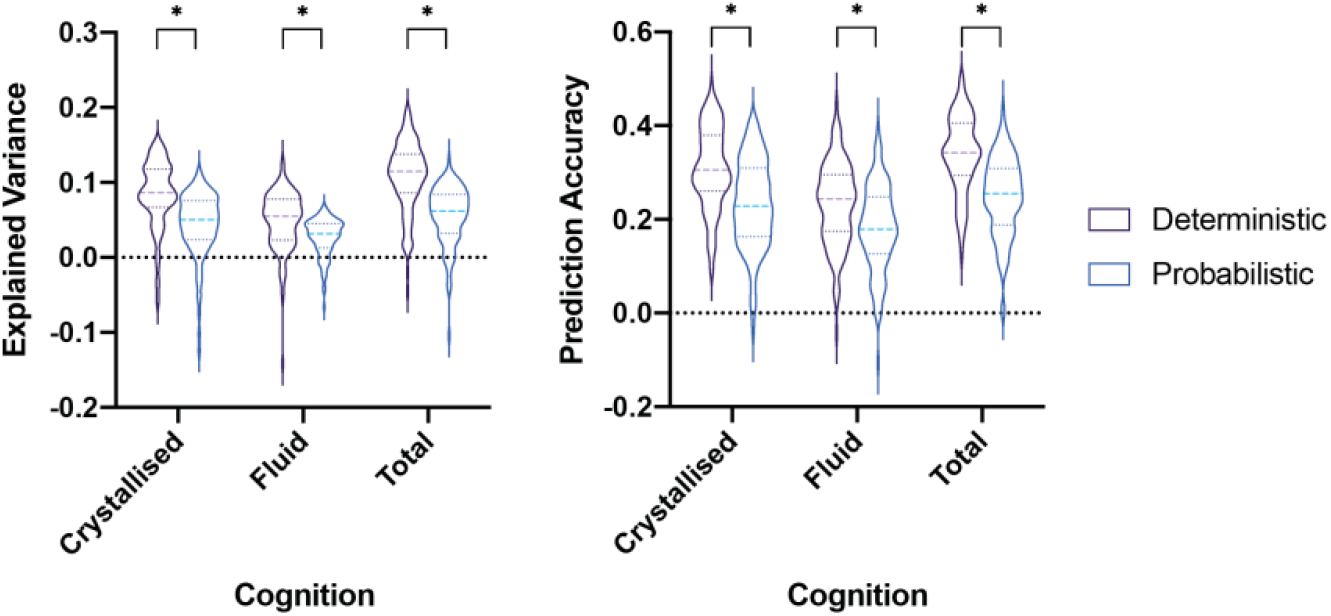
Deterministic versus Probabilistic Tractography. Explained variance (left) and prediction accuracy (right) violin plots for structural connectivity derived from deterministic and probabilistic tractography to predict crystallised, fluid and total cognition. Solid lines indicate the distribution of values, dashed lines indicate the median, and dotted lines indicate the interquartile range. Pairwise differences between models to predict each cognition composite using structural connectivity derived from deterministic and probabilistic tractography were evaluated using a Mann Whitney U Test. * indicates significant differences (p < 0.05) after Benjamini-Hochberg correction for multiple comparisons.

## Discussion

In this study, we evaluated whether integrating functional and structural connectivity improves predictions of cognitive scores in 415 healthy young adults. Using whole brain resting-state functional connectivity, structural connectivity, hybrid function-structure connectivity, and ensemble predictions, we predicted individual crystallised, fluid, and total cognition. The conclusions of the study are three-fold. First, we demonstrate that integrating functional and structural connectivity information at both an input level, via hybrid connectivity, and an output level, via ensemble predictions, modestly but significantly outperforms functional and structural connectivity alone to predict individual crystallised, fluid, and total cognitive abilities. Second, we show that distinct functional and structural connections are important to predict crystallised, fluid, and total cognition. Lastly, we find that structural connectivity derived from deterministic tractography significantly outperforms structural connectivity derived from probabilistic tractography to predict individual cognitive scores.

Prior studies have implemented machine (He et al., 2020, Kong et al., 2018, Li et al., 2019) and deep learning (He et al., 2020) algorithms to predict behaviour and cognition in the Human Connectome Project dataset using functional connectivity. Li et al. reported cross-validated prediction accuracies ranging from approximately 0.1 to 0.4 to predict individual scores on the Dimensional Change Card Sort, Flanker Inhibitory Control and Attention, Picture Sequence Memory, Picture Vocabulary, Oral Reading Recognition, List Sorting Working Memory, and Pattern Comparison Processing Speed tasks using FC (Li et al., 2019). He et al. demonstrated that kernel regression, fully connected neural networks, and BrainNetCNN (Kawahara et al., 2017) achieve comparable prediction accuracies when used to predict behavioural and cognitive measures using FC (He et al., 2020). They reported prediction accuracies ranging from approximately 0.1 to 0.4 to predict individual scores on the Dimensional Change Card Sort, Flanker Inhibitory Control and Attention, Picture Sequence Memory, Picture Vocabulary, Oral Reading Recognition, List Sorting Working Memory, and Pattern Comparison Processing Speed tasks (He et al., 2020). While we do not predict scores on these individual tasks (as they are noisier and may be less reliable than the composites), these scores are used to derive the Crystallised, Fluid, and Total cognition composite.

White matter pathways and neural co-activation patterns in the brain produce complex cognitive functions. While brain functional and structural connections are undeniably related, most studies analyse their independent contributions to behaviour. Here, we replicate prior findings that functional and structural connectivity can be used separately to predict crystallised and fluid cognition. More importantly, we demonstrate that integrating functional and structural connectivity significantly improves cognitive predictions. We show that while functional connectivity can predict the composite scores, structural connectivity can also predict the composite scores with comparable explained variance and prediction accuracy. We also demonstrate that integrating functional and structural connectivity at both an input level via hybrid connectivity and at an output level via ensembling significantly improves the explained variance and prediction accuracies. This suggests that functional and structural connectivity capture unique and complementary information, and when combined can help us better understand the neural correlates of cognition.

Zimmerman et al. examined how functional and structural connectivity independently map to cognition (Zimmermann et al., 2018); however, they did not integrate the two modalities to quantify and compare their joint contribution to cognitive scores. They report that a small number of distributed structural connections and a larger set of cortical and cortico-subcortical functional connections show relationships with cognition and conclude that functional and structural connections uniquely map onto cognition (Zimmermann et al., 2018). Our results support this conclusion: specifically, we found that structural and functional connections linking cortical areas in the frontal, temporal, parietal, and cingulate cortices with subcortical regions such as the caudate are strongly predictive of crystallised and fluid abilities. We quantify the lack of correlation between pairwise feature importance for functional and structural connections, and between regional feature importance from models using functional and structural connectivity to predict crystallised and fluid cognition. We also demonstrate that distinct functional and structural connections are important to predict crystallised and fluid cognition (described below). These results are in line with Zimmerman et al.’s (Zimmermann et al., 2018) findings that unique functional and structural connections and regions may underlie different aspects of cognitive function. We argue here that both functional and structural connections should be integrated to fully understand the neural correlates of cognition.

Amico and Goni (Amico and Goñi, 2018) simultaneously studied functional and structural connectivity patterns related to changes in brain networks across seven tasks (gambling, relational, social, working memory, motor, language, emotion) and resting-state. The connICA methodology extracts robust functional connectivity patterns from a set of functional connectomes using independent component analysis (ICA) (Amico et al., 2017). This technique was extended to include both structural and functional connectomes by merging them into a “hybrid” matrix (Amico and Goñi, 2018). This hybrid connICA approach disentangled two task-dependent components: one encompassing within- and between- network connections of dorsal attentional and visual structures, as well as subcortical areas, and a second including connectivity between visual, frontoparietal, default mode, and subcortical networks (Amico and Goñi, 2018). Amico and Goni showed that by combining functional and structural connectivity, meaningful information can be extracted from heterogeneous brain networks while capturing individual differences (Amico and Goñi, 2018). In this study, we adopted a modified version of this approach to study the relationship between hybrid connectivity and cognitive functioning. We integrated functional and structural information at either the model input or output level. At the input level, we concatenated functional and structural connectivity to generate hybrid connectivity. At the output level, we averaged the predictions of the cognitive scores from models created separately using functional and structural connectivity to generate ensemble predictions. Prior work has shown that the use of ensemble predictions integrating information across parcellations in neuroimaging data can be used to improve sex (Dhamala et al., 2019) and disease (Khosla et al., 2019) classification. Here, we show that ensemble predictions integrating information about imaging modalities can also significantly improve prediction of cognitive abilities.

Cattell and Horn’s two-component theory of intellectual development proposes a distinction between crystallised and fluid abilities in how they develop and transform throughout life (Cattell, 1967, Horn and Cattell, 1967, Horn and Cattell, 1966). Crystallised intelligence is the ability to use learned knowledge, experience, and skills, and fluid intelligence is the ability to solve new problems using logic, encode new episodic memories, and adapt to novel situations in everyday life (Heaton et al., 2014). In the HCP dataset (Van Essen et al., 2013), the NIH Toolbox Cognition Battery was used to assess crystallised and fluid abilities. Crystallised abilities are influenced by education and cultural factor, and fluid abilities, while also dependent on educational and cultural factors, are more dependent on biological processes within neural structures that enable brain function (Cattell, 1967, Horn and Cattell, 1967, Horn and Cattell, 1966, Heaton et al., 2014). Interestingly, in our results functional and structural connectivity patterns were less predictive of fluid abilities than of crystallised abilities. In the NIH Toolbox Cognition Battery, the Crystallised Cognition Composite reflects scores from tasks measuring vocabulary and reading decoding, while the Fluid Cognition Composite reflects scores from tasks measuring cognitive flexibility, inhibitory control and attention, episodic memory, working memory, and processing speed. The eloquent nature of the mapping between brain anatomy/physiology and language, including vocabulary and reading as measured by the Crystallised Cognition Composite, may explain the increase in prediction accuracy of those scores when compared to the Fluid Cognition Composite that may rely on several overlapping brain networks. Another possible explanation for the higher predictability of crystallised abilities (relative to fluid) lies in the impact of environment on the brain’s connectomes. Functional and structural connectivity have been shown to be related to learning and life experience (Tooley et al., 2019, Zatorre et al., 2012, Peng et al., 2018, Johansen-Berg et al., 2010). Hence, it is possible that the joint impact of environment on connectivity networks and crystallised abilities means it is easier to predict one from the other.

Many functional connections are negatively related to individual cognitive abilities as shown in Figure 3. These results align with those previously reported by Zimmerman et al. (Zimmermann et al., 2018): a large number of interhemispheric cortico-cortical connections negatively map onto an array of cognitive measures. Negative feature importance for functional connections can be interpreted in two ways: it can reflect an inverse relationship with positive functional correlations; or it can reflect a direct relationship with negative functional correlations. However, anticorrelations, or negative functional connections, account for a mere 6.0% of connections prior to global signal regression across all subjects (including only 1.8% < −0.05). Hence, it is unlikely that these anticorrelations strongly contribute to predictions of cognitive performance.

Crystallised cognition, as measured by the NIH toolbox, mainly represents language (vocabulary and reading decoding) abilities. Here, we identify, unsurprisingly, that functional and structural connectivity with brain regions involved in language and auditory processing are especially important to predict crystallised cognition. Specifically, we find that connections involving the pars opercularis, which is part of Broca’s area, temporal pole, banks of the superior temporal sulcus, supramarginal, and superior temporal areas are important. Moreover, we also observe that regions in the frontal lobe responsible for higher order cognitive processes are important. Regions of the middle and superior prefrontal cortex are implicated in verbal working memory and retrieval of words from semantic memory (Nyberg et al., 2003, Lee et al., 2000), which may explain the importance of prefrontal regions in our models.

Fluid cognition represents a wide range of cognitive processes: executive function (cognitive flexibility and inhibitory control and attention), episodic memory, working memory, and processing speed. These processes each rely on distributed connections through the brain. In our results, we show that functional and structural connections important to predict fluid cognition are dispersed throughout the cortex and subcortex. As shown in Figures 3 and 5, cortico-subcortical connections involving the caudate, putamen, thalamus, and amygdala with regions of the frontal, parietal, and temporal lobes emerge as predictive of fluid cognition. The cortical regions implicated in fluid cognition comprise nodes of the frontoparietal network, dorsal attention network, cingulo-opercular task control network, and default mode network, all of which have been implicated in executive function (Reineberg et al., 2015, Spreng et al., 2010, Leech et al., 2011, Jaywant et al., 2020) and memory (Iidaka et al., 2006, Wallis et al., 2015, Dixon et al., 2017). The frontoparietal, dorsal attention, and cingulo-opercular networks are frequently implicated in maintaining and updating information in mind, shifting attention, inhibiting distractions, and facilitating goal-directed behaviour (Spreng et al., 2010, Iidaka et al., 2006, Wallis et al., 2015, Dixon et al., 2017). The default mode network underlies self-referential thinking and future simulation and has been shown to be related to episodic memory (Fair et al., 2008, Sheline et al., 2009, Spreng and Grady, 2010). Connections between cortical regions and the hippocampus were also implicated in fluid cognition, likely because the composite measure includes tasks assessing episodic memory. More broadly, our findings suggest that more distributed functional connections are relevant to a heterogenous set of cognitive functions that underlie adaptive, goal-directed behaviour in novel situations (i.e. fluid cognition). In other words, if fluid cognition helps us plan and act in novel situations and solve problems, it is likely going to be dependent on a wide variety of cognitive skills (and brain regions) than acquired knowledge.

The caudate nucleus is fundamental to successful goal-directed action: it excites correct action schemas and selects appropriate sub-goals by evaluating action-outcomes (Grahn et al., 2008). Here, we observe that structural connections between the caudate and regions in frontal, parietal, occipital, and cingulate cortices are especially important to predict crystallised and fluid cognition. This finding is consistent with the known role of corticostriatal-thalamocortical loops in healthy executive function (Seger, 2009), and in neurologic disorders (Shepherd, 2013, Leisman et al., 2013) that result in executive dysfunction such as Parkinson’s disease (Zgaljardic et al., 2006) and Huntington’s disease (Rangel-Barajas and Rebec, 2016).

In our results, we observe that many distributed cortical and subcortical regions are equally important when using SC to predict cognition, but only a few regions are important when using FC to predict cognition. Previous work (Jaywant et al., 2020) using a partial least squares regression approach to identify the relationship between inferred structural disconnection and cognitive inhibition observed that it was mostly structural connections that were important to predict cognition, with only two functional connections contributing additional unique variance. Our results here align with those findings; While functional and structural connections uniquely predict cognitive abilities, at a regional level there are more regions that are important for the prediction when using structural connectivity than when using functional connectivity. That is, there are a few regions whose functional connectivity profile as a whole largely outweigh the other regions’ contributions to prediction; however, this is not the case for regional structural connectivity profiles.

Myelinated axons form short and long white matter pathways which comprise neural circuits involving cortical and subcortical regions of the brain. These pathways, the communication highways of our brain, can be estimated from dMRI using tractography algorithms, which generally follow these three steps: 1) estimate voxel-wise local fibre orientations, 2) link local fibre orientations to generate streamline trajectories of white matter fibres, and 3) identify which grey matter regions the streamlines connect and generate a connectivity matrix (Sarwar et al., 2019). Two main classes of tractography algorithms, deterministic and probabilistic, differ in how they sample fibre directions for streamline propagation: fixed orientations direct streamlines in deterministic algorithms, while probabilistic algorithms estimate a distribution of fibre orientations at each voxel and a sample is randomly drawn from this distribution to direct the streamline propagation (Sarwar et al., 2019). Deterministic approaches cannot account for inherent uncertainty in fibre orientation estimates and are sensitive to the principal direction(s) estimated (Sarwar et al., 2019). Probabilistic approaches overcome this issue but are more computationally expensive (Sarwar et al., 2019). Prior work has found that probabilistic tractography more faithfully reconstructs connectome phantoms when fibre complexity exceeds that of in vivo diffusion MRI data, but falls short of deterministic tractography when mapping lower complexity phantoms, comparable to in vivo data, due to an abundance of false-positive connections (Sarwar et al., 2019). In this study, we find that structural connectivity derived from deterministic tractography significantly outperforms structural connectivity derived from probabilistic tractography to predict individual crystallised and fluid abilities.

## Limitations

Machine learning algorithms using neuroimaging data are prone to the curse of dimensionality. Voxel-wise imaging data, on the order of hundreds of thousands of features, and regional data can have hundreds or thousands of features. Here, we parcellate the brain into 86 regions: 34 cortical regions and 9 subcortical regions per hemisphere ^30^. When taking the upper or lower triangular of the pairwise functional and structural connectivity matrices, this leaves us with 3655 features. Dimensionality reduction through parcellation decreases noise, reduces computational cost, and enables more interpretable models, but loses valuable information captured in the voxel-wise data; Future work performing a voxel-wise analysis of the data can address this issue.

In this study, we only used data from the Human Connectome Project. Although we exclusively evaluate our models on test sets not used to train the models and perform 100 permutations of each model with unique train/test splits, the results we report here may not be generalisable to other datasets; Future work performing out-of-dataset evaluations can address this limitation.

We show that many regions are equally important when using SC to predict cognition, but only a few regions are important when using FC to predict cognition. While interesting, these results may be due to the differences in distribution of FC and SC entries and the optimised model hyperparameters using FC and SC. We resampled the SC matrices using a Gaussian distribution with a mean of 0.5. Meanwhile, the FC matrices, which were global signal regressed and z-transformed, have a Gaussian distribution with a mean of 0. Although the variables are normalised within the machine learning pipeline, these differences in FC and SC may underlie the observed differences in regional importance.

Age (Damoiseaux, 2017, Song et al., 2014), sex (Gong et al., 2011, Gur and Gur, 2017, Satterthwaite et al., 2015, Weis et al., 2019, Ingalhalikar et al., 2014, Jacobs and Goldstein, 2018, Jacobs et al., 2016), and environment/experience (Tooley et al., 2019, Sripada et al., 2014) influence connectivity. Hence, it is likely that they, along with other demographic variables such as gender and ethnicity, may influence the relationship between connectomics and cognition (Jiang et al., 2020). Future work should examine how the relationship between connectomics and cognition varies based on demographics.

## Conclusion

Understanding neural correlates of cognition in healthy individuals is a critical first step towards understanding changes in cognitive functioning as a result of age, injury, and disease. Here, we integrate neuroimaging, connectomics, and machine learning approaches to explore how multimodal brain connectivity predicts individual crystallised and fluid abilities. We report three main findings. First, integrating functional and structural connectivity significantly outperforms the independent use of functional connectivity and structural connectivity to predict individual crystallised and fluid cognition. Second, distinct functional and structural connections are important to predict crystallised and fluid cognition. Third, structural connectivity derived from deterministic tractography significantly outperforms structural connectivity derived from probabilistic tractography to predict crystallised and fluid cognition. Taken together, this suggests the integration of multimodal connectivity is crucial to understand the neuroanatomical and neurophysiological correlates of cognition.

## Citation Gender Diversity Statement

Recent work in neuroscience and other fields has identified a bias in citation practices such that papers from women and other minorities are under-cited relative to the number of such papers in the field (Maliniak et al., 2013, Caplar et al., 2017, Chakravartty et al., 2018, Thiem et al., 2018, Dion et al., 2018, Dworkin et al., 2020). Here we sought to proactively consider choosing references that reflect the diversity of the field in thought, form of contribution, gender, and other factors. We used classification of gender based on the first names of the first and last authors (Dworkin et al., 2020), with possible combinations including male/male, male/female, female/male, and female/female. Excluding self-citations to the first and last authors of our current paper, the references contain 54.7% male/male, 17.4% male/female, 19.8% female/male, and 8.1% female/female. We look forward to future work that could help us to better understand how to support equitable practices in science.

## Author Contributions

Conceptualization, E.D. and A.K.; Methodology, E.D. K.W.J. and A.K.; Software, E.D., S.D; Investigation, E.D., S.D.; Formal Analysis, E.D.; Resources, A.K; Data Curation, E.D. and K.W.J.; Writing – Original Draft, E.D.; Writing – Review & Editing, E.D., K.W.J., A.J., S.D., A.K.; Visualisation, E.D.; Supervision, A.K.; Funding Acquisition, A.K.

## Acknowledgements

Data were provided and made available by the Human Connectome Project, WU-Minn Consortium (Principal Investigators: David Van Essen and Kamil Ugurbil; 1U54MH091657) funded by the 16 NIH Institutes and Centres that support the NIH Blueprint for Neuroscience Research; and by the McDonnell Centre for Systems Neuroscience at Washington University. Data can be accessed at https://www.humanconnectome.org/study/hcp-young-adult/document/1200-subjects-data-release.

The authors would like to acknowledge Dr. Mallar Chakravarty from McGill University for contributions to an earlier version of this work and insightful comments on the manuscript. The authors would also like to thank Raihaan Patel from McGill University for contributions to an earlier version of this work.

This work was supported by the following National Institutes of Health (NIH) grants: R21 NS104634-01 (AK) and R01 NS102646-01A1 (AK). The sponsor did not have any role in the study design, the analysis, or interpretation of the data; the writing of the report; or the decision to submit the manuscript for publication.

## Data and Code Availability Statement

The data that support the findings of this study are openly available as part of the Human Connectome Project at https://www.humanconnectome.org/study/hcp-young-adult/document/1200-subjects-data-release.

The codes generated during this study are available on GitHub (https://github.com/elvisha/CognitivePredictions).

