## Supplemental Materials for "Integrating multimodal connectivity improves prediction of individual cognitive abilities"

Regional positive and negative feature importance were computed by taking the sum over that region’s pairwise positive and pairwise negative feature importances. There are no correlations between regional feature importance for any of the following pairs: 1) positive functional and positive structural regional feature importance in the crystallised abilities model, 2) negative functional and negative structural regional feature importance in the crystallised abilities model, 3) positive functional and positive structural regional feature importance in the fluid abilities model, 4) negative functional and negative structural regional feature importance in the fluid abilities model, 5) positive functional and positive structural regional feature importance in the total cognitive abilities model, 6) negative functional and negative structural regional feature importance in the total cognitive abilities model. This further strengthens our results presented in the main paper that distinct functional and structural connectivity information is important to predict cognition. Two regions where the functional and structural regional importance are particularly different are the left caudal middle frontal and the left supramarginal gyrus, as shown in Supplementary Figure 1.

Supplementary Figure 1: Regional positive and negative feature importance.

Regional positive and negative feature importance from models using functional connectivity (left) and structural connectivity (right) to predict Crystallised (top), Fluid (middle), and Total (bottom) Cognition Composites.

Regional positive feature importance reflects the sum of all positive feature weights for that region; regional negative feature importance reflects the sum of all negative feature weights.

Lateral and medial views of the right (RH) and left (LH) hemispheres are shown.
